# An advanced strategy for comprehensive profiling of ADP-ribosylation sites using mass spectrometry-based proteomics

**DOI:** 10.1101/501353

**Authors:** Ivo A. Hendriks, Sara C. Larsen, Michael L. Nielsen

**Affiliations:** Proteomics program, Novo Nordisk Foundation Center for Protein Research, Faculty of Health and Medical Sciences, University of Copenhagen, Blegdamsvej 3B, 2200 Copenhagen, Denmark

**Author notes:** These authors contributed equally (Ivo A. Hendriks, Sara C. Larsen).

## Abstract

ADP-ribosylation is a widespread post-translational modification (PTM) with crucial functions in many cellular processes. Here, we describe an in-depth ADP-ribosylome using our Af1521-based proteomics methodology for comprehensive profiling of ADP-ribosylation sites, by systematically assessing complementary proteolytic digestions and precursor fragmentation through application of electron-transfer higher-energy collisional dissociation (EThcD) and electron transfer dissociation (ETD), respectively. While ETD spectra yielded higher identification scores, EThcD generally proved superior to ETD in identification and localization of ADP-ribosylation sites regardless of protease employed. Notwithstanding, the propensities of complementary proteases and fragmentation methods expanded the detectable repertoire of ADP-ribosylation to an unprecedented depth. This system-wide profiling of the ADP-ribosylome in HeLa cells subjected to DNA damage uncovered >11,000 unique ADP-ribosylated peptides mapping to >7,000 ADP-ribosylation sites, in total modifying over one-third of the human nuclear proteome and highlighting the vast scope of this PTM. High-resolution MS/MS spectra enabled identification of dozens of proteins concomitantly modified by ADP-ribosylation and phosphorylation, revealing a considerable degree of crosstalk on histones. ADP-ribosylation was confidently localized to various amino acid residue types, including less abundantly modified residues, with hundreds of ADP-ribosylation sites pinpointed on histidine, arginine, and tyrosine residues. Functional enrichment analysis suggested modification of these specific residue types is directed in a spatial manner, with tyrosine ADP-ribosylation linked to the ribosome, arginine ADP-ribosylation linked to the endoplasmic reticulum, and histidine ADP-ribosylation linked to the mitochondrion.

## INTRODUCTION

ADP-ribosylation is an emerging reversible post-translational modification (PTM) catalyzed by a large and diversified group of enzymes known as ADP-ribosyltransferases (ARTs). Based on their conserved structural features of the active center, the enzymes can be divided into two major groups: ARTCs (cholera toxin-like) and ARTDs (diphtheria toxin-like, also known as poly(ADP-ribose)polymerases (PARPs)^1^. Of these, the ARTCs all catalyze mono-ADP-ribosylation (MARylation), where one ADP-ribose moiety is transferred from NAD+ to the target protein. The PARPs/ARTDs constitute a larger group, with members able to catalyze either MARylation, or poly-ADP-ribosylation (PARylation), where one or more ADP-ribose moieties are transferred to an already protein-bound ADP-ribose. Especially the role of nuclear PARylation catalyzed by the PARP1 and PARP2 enzymes during the DNA damage response has been comprehensively studied, and has led to the development and approval of PARP inhibitors for treatment of ovarian and breast cancer^2–5^.

Similar to most PTMs, the cellular abundance of ADP-ribosylation is very low, and further complicated by the rapid turnover of the modification *in vivo*^6^, making an enrichment step prior to mass spectrometric (MS) analysis necessary. An additional challenge for studying site-specific ADP-ribosylation is the lability of the peptide-bound ADP-ribose group compared to the peptide amine bond. This, combined with the many reported amino acid acceptor sites^7–17^, renders confident localization of ADP-ribosylation analytically challenging when using conventional MS approaches. Several strategies have been explored for enrichment of ADP-ribosylation, with each possessing distinct advantages and disadvantages. One method combines boronate-affinity chromatographic enrichment of ADP-ribosylated peptides with elution using hydroxylamine, generating a hydroxamic acid derivative on glutamic acid and aspartic acid residues with a distinctive mass of 15.0109 Da^17^. While this method is capable of identifying a high number of sites, the ADP-ribosylation sites reported were exclusively observed under non-physiological conditions using cellular knock-down (siRNA) of the PAR-degrading glycohydrolase PARG^17^. Moreover, a major disadvantage of this approach is the analytical bias towards aspartic acid and glutamic acid residues, since other acceptor residues cannot be identified. An alternative method makes use of phosphodiesterases derived from snake venom, to hydrolyze PAR into a phosphoribose remnant suitable for enrichment using well-established phosphorylation proteomics approaches^18^. Similarly, in a recently published method, hydrofluoric acid is used to convert ADP-ribose into ribose adducts^19^. While all of these methods are able to report ADP-ribosylation on various amino acid residues, they commonly lack sensitivity for in-depth characterization of the ADP-ribosylome, especially for investigation of ADP-ribosylation under physiological conditions. Besides, a major disadvantage of the methods described is that none of them identifies the actual peptide-bound ADP-ribose group, in contrary to most PTM-based methodologies where the original PTM remains bound to the analyzed peptide and therefore directly detectable by MS analysis^20,21^.

To mitigate these challenges, we have previously demonstrated the use of the macrodomain Af1521 for comprehensive and specific enrichment of ADP-ribosylation sites^22–25^. While this methodology allows for enrichment of the attached and intact ADP-ribose, the lability of ADP-ribosylation still constitutes an analytical challenge for site-specific identification when using MS-based analysis utilizing higher-energy collisional dissociation (HCD) fragmentation^22^. Conversely, the non-ergodic fragmentation propensity of electron transfer dissociation (ETD) fragmentation has proven superior for identification and confident localization of labile PTMs including phosphorylation^26,27^, and we have also previously confirmed this for ADP-ribosylation^22^.

However, one limitation of ETD fragmentation is the generation of dominant charge-reduced precursor ions, thus providing poorer fragmentation efficiency, and a bias against peptides with lower charge states^28^. Several strategies has been explored to overcome the limitations of ETD fragmentation, including the use of alternative proteases^29^. In shotgun proteomics trypsin is the most widely used protease due to the specific and efficient cleavage C-terminal to arginine and lysine residues^30^, generating peptides of an average length that is convenient for mass spectrometric resolution^31^. Digesting with endoproteinase Lys-C generates longer peptides on average carrying higher charge states, making them more suitable for analysis using ETD fragmentation^32^. Additionally, a combination of complementary fragmentation techniques, ETD and HCD, referred to as electron-transfer higher-energy collisional dissociation (EThcD), helps to overcome the challenge of unreacted and charge-reduced precursors that are often highly abundant upon ETD fragmentation, while concomitantly obtaining b- and y-ions alongside an increase in c- and z-ion formation^33^. Notably, EThcD has proven superior for confident localization of labile PTMs including phosphorylation^34^ and glycosylation^35^, and has previously been used for identification of ADP-ribosylated peptides^36^.

Here, we evaluated the usefulness of performing sample digestion with either Lys-C or trypsin prior to enrichment of ADP-ribosylated peptides using an augmented version of our Af1521 enrichment strategy, as well as the potential of ETD and EThcD fragmentation for localizing the ADP-ribose to the correct amino acid residue. Collectively, we describe an improved analytical MS-based proteomics strategy for enrichment, identification, and localization of ADP-ribosylation sites at the systems-wide level. By combining different proteases with various precursor fragmentation strategies, we report confident identification of >7,000 ADP-ribosylation sites (localization probability >0.90) in cultured human cells subjected to DNA damage, representing a comprehensive profile of the human ADP-ribosylome.

## RESULTS

To enable comprehensive and robust profiling of ADP-ribosylation, we initially set out to optimize our established proteomics methodology. To this end, we relied on our peptide-level enrichment strategy, which includes reduction of poly-ADP-ribose (PAR) to monomers (MAR) using the PAR glycohydrolase (PARG) enzyme, followed by purification of the ADP-ribosylated peptides using the Af1521 macrodomain^22,25^. Importantly, we sought to optimize the protocol to make it experimentally more robust, and achieve the highest degree of peptide purity (Figure 1A). Key improvements include a hydrophilic purification of the protein digest, followed by lyophilization to maximize recovery of peptides that harbor ADP-ribosylation while eliminating large hydrophobic molecules. Following PARG treatment of the samples and subsequent ADP-ribosylation enrichment using Af1521, we incorporated a size-exclusion filtering step to remove the Af1521 macrodomain from the elutions, which is the major contaminant in previous Af1521 experiments^25^. Finally, high-pH fractionation of the remaining peptides reduced sample complexity, and moreover served as a final purification step owing to the unique hydrophilic nature of ADP-ribosylated peptides (detailed in Methods section).

**Figure 1.**
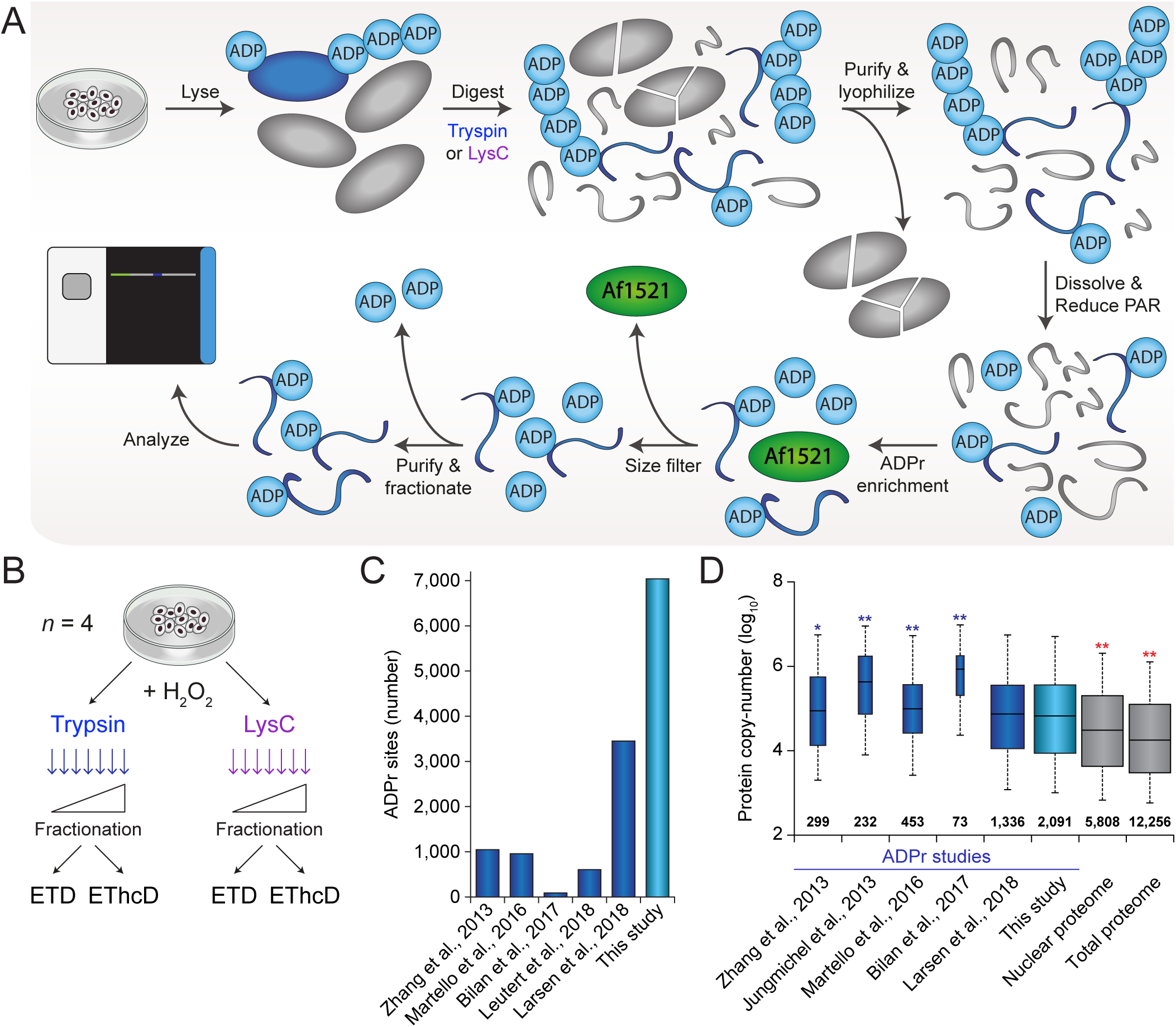
Optimized proteomics methodology for deep profiling of the ADP-ribosylome. (A) Overview of the general experimental workflow used for purification of ADP-ribosylated peptides. Sample preparation starts with lysis of the starting material, which was cultured human cells, but can in principle also be tissue material or samples derived from other model organisms. Lysis and homogenization is performed in guanidine buffer, to denature all proteins and preserve labile PTMs, and followed by digestion of the proteins by either trypsin or Lys-C. Subsequently, hydrophilic peptides are purified through reversed-phase chromatography, and salts as well as larger particulates such as partially digested proteins and other contaminants are excluded at this stage. Purified peptides are lyophilized to maximize their recovery, and dissolved in a mild buffer to facilitate efficient reduction of ADP-ribose polymers with the PARG enzyme. The same buffer enables efficient enrichment of ADP-ribosylated peptides using the Af1521 macrodomain, which is afterwards removed using a size-exclusion filtration step that removes large polypeptides. The final step in the sample preparation is clean-up using StageTips, optionally with on-StageTip high-pH reversed-phase fractionation to reduce sample complexity. Purified ADP-ribosylated peptides may then be analyzed using mass spectrometry, ideally using ETD-type precursor fragmentation to correctly localize the ADP-ribose. (B) The experimental design for this study. Cell cultures were performed in quadruplicate; *n*=4, and all cells were treated with H_2_O_2_ to induce ADP-ribosylation. (C) Overview of the total number of sites identified in this study, as well as the numbers reported by previous ADP-ribosylation proteomics studies^17,22,23,25,36^. Note that these numbers are irrespective of the different purification methods and bioinformatics approaches used. (D) Depth of sequencing analysis, inferred through known copy-numbers for proteins identified to be ADP-ribosylated in this study, and in various other studies^17,22,25,36,51^. Protein copy-numbers were derived from a recent deep HeLa proteome study^37^, which covers the vast majority (>99%) of all ADP-ribosylated proteins identified. The width of the boxplots (for ADP-ribosylation studies) and the numbers set above the horizontal axis correspond to the number of proteins identified. Whiskers; 95^th^ and 5^th^ percentiles, box limits; 3^rd^ and 1^st^ quartiles, center bar; median. Significance of differences between depth of sequencing was assessed by Student’s two-tailed *t*-testing, blue and red asterisks indicate that our study achieved more or less depth, respectively. * *p*<0.05, ** *p*<0.001.

To achieve a greater depth and width of ADP-ribosylation sequencing, we expanded our core experimental workflow in two ways (Figure 1B). First, we performed initial digestion of proteins either using trypsin or the endoproteinase Lys-C. Unlike trypsin, Lys-C does not cleave C-terminal of arginines, thus resulting in a different peptide composition. Second, samples were analyzed using either electron-transfer disassociation (ETD), or using ETD with supplemental higher-collisional dissociation (EThcD). In all cases, experiments were carried out using quadruplicate HeLa cell cultures, which were all subjected to treatment with hydrogen peroxide, and all samples were high-pH fractionated prior to analysis, as described previously^22^.

Prior to analyzing our main samples, we set out to investigate a suitable level of supplemental activation energy for EThcD fragmentation of ADP-ribose-modified peptides. To this end, we enriched ADP-ribosylated peptides from HeLa cells either mock treated or treated with hydrogen peroxide, using our established workflow (Figure S1A). Following purification, samples were split in half, and analyzed back-to-back using either pure ETD or EThcD at increasing levels of supplemental activation energy (Figure S1B). Overall, we observed an increasing propensity for EThcD to successfully localize ADP-ribosylation on peptides when using higher energies. However, when increasing energy over 20, peptide scores decreased and fewer ADP-ribose-modified peptides could be identified. Thus, EThcD with a supplemental activation setting of 20 was used for all main experiments.

### In-depth profiling of the human ADP-ribosylome

All ADP-ribosylation samples were analyzed using an Orbitrap Fusion Lumos Tribrid mass spectrometer with either ETD or EThcD fragmentation (Figure 1A-B), and all data was processed in a single computational run using semi-restricted search settings^22^. Importantly, our experimental methodology in combination with ETD-driven peptide fragmentation allows for unbiased and accurate assignment of the ADP-ribose group to any of the nine reactive amino acid residues that have been reported previously.

In total, we identified 11,265 unique ADP-ribose-modified peptides (Table S1), mapping to 7,040 unique ADP-ribosylation sites (Table S2), across 2,149 human proteins (Figure 1C and Table S3). We assessed the depth of sequencing by linking identified ADP-ribosylation target proteins with established protein copy-numbers^37^, and found that our human ADP-ribosylome is significantly deeper than most previously published studies (Figure 1D). Although we did not significantly increase depth of sequencing compared to our most recent study^22^, we nonetheless increased the number of identified ADP-ribosylation sites and ADP-ribosylated proteins by ~100% and ~50%, respectively. The dynamic range of ADP-ribosylated protein intensities measured across our experiments spanned nearly six orders of magnitude (Figure S1C), in line with most recent deep proteome studies, and overall highlighting the analytical challenge of proteomic analysis of PTMs, which often populate a very large dynamic range. With the majority of ADP-ribosylated target proteins residing within the nucleus (Figure S1F), our data collectively outlines that ADP-ribosylation can modify over one-third of all known nuclear proteins.

Both enzymatic and fragmentation approaches proved highly reproducible (Figure 2A), and succeeded in identifying and localizing thousands of ADP-ribosylated peptides. In general, MS/MS spectra resulting from pure ETD fragmentation had higher Andromeda scores and delta scores (Figure S2A), with EThcD more frequently identifying peptides with higher mass and higher mass-over-charge (m/z)(Figures S2B-C). Outside of peptide mass, these properties were not significantly different between peptides derived from trypsin or Lys-C digestion. However, when considering peptides uniquely identified by either pure ETD or EThcD (Figure S2D), we found that EThcD outperformed ETD.

**Figure 2.**
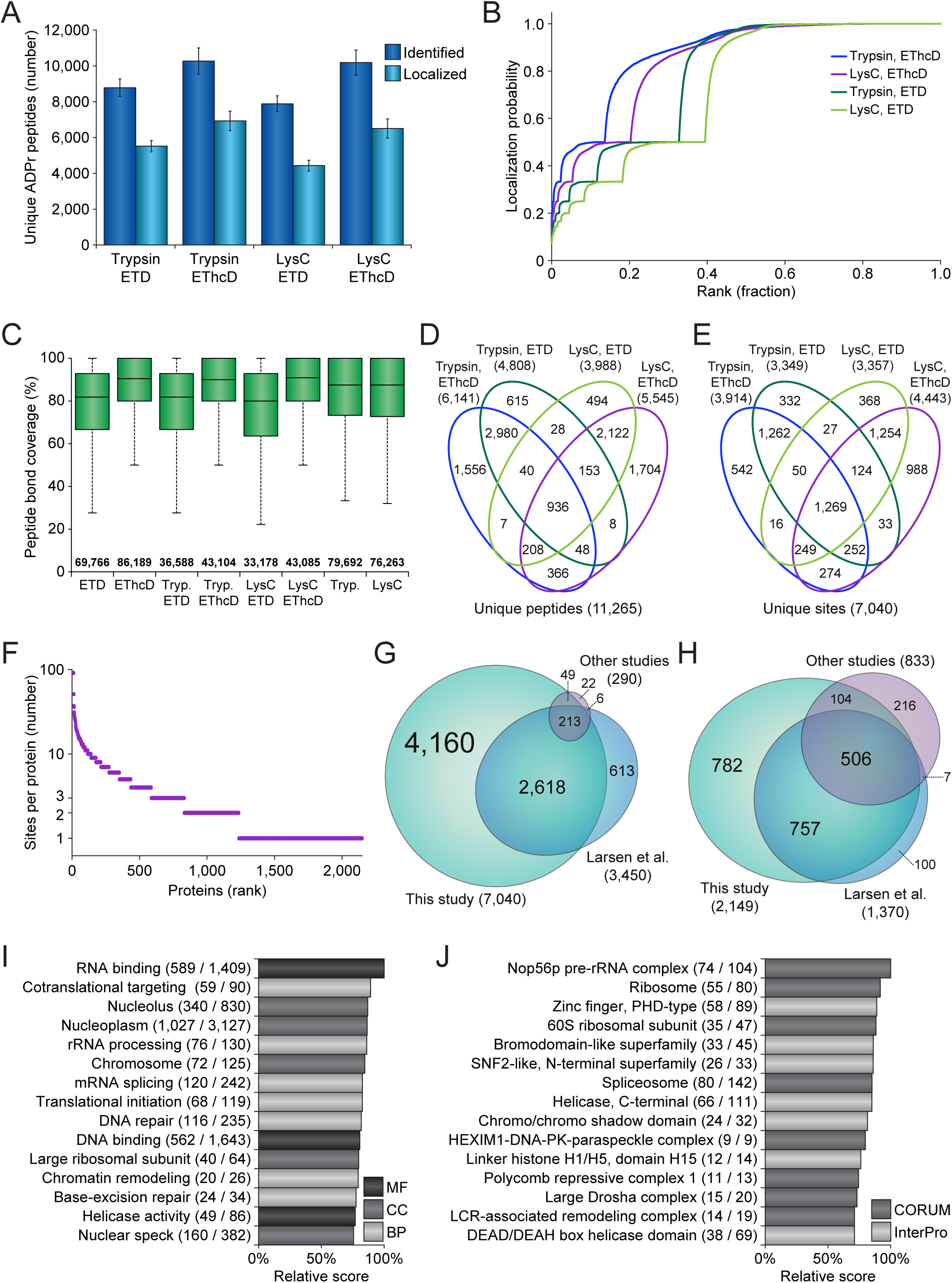
In-depth profiling of the human ADP-ribosylome. (A) Overview of the number of unique ADP-ribosylated peptides identified and localized across the four different experimental conditions. Error bars represent SD, *n*=4 cell culture replicates. (B) ADP-ribose localization probability plotted against the ranked fraction of all peptide-spectrum-matches (PSMs) resulting from the four different approaches. Note that although all probabilities are displayed, only those over 0.9 were used for assignment of ADP-ribosylated peptides and sites. (C) Peptide bond coverage analysis, which demonstrates the fraction of the maximum number of peptide bonds for which peptide bonds were fragmented, and for which peaks were detected and assigned. Whiskers; 1.5× interquartile range (IQR), box limits; 3^rd^ and 1^st^ quartiles, center bar; median. Numbers set above the horizontal axis correspond to the number of identified ADP-ribosylation PSMs. (D) Venn diagram depicting the distribution of unique ADP-ribosylated peptides identified across the four different approaches. (E) As **D**, but for unique ADP-ribosylation sites. (F) Overview of the number of ADP-ribosylation sites per protein, plotted against the total number of identified proteins ranked by site number. (G) Scaled Venn diagram visualizing distribution of identified ADP-ribosylation sites in this study compared to other ADP-ribosylation studies^16,22,36^. “Other studies”; Bilan et al., and Bonfiglio et al. For this comparison, only studies using unbiased purification methods, and unrestricted or semi-restricted database searches, were considered. (H) Scaled Venn diagram visualizing distribution of identified ADP-ribosylation target proteins in this study, compared to other ADP-ribosylation proteomics studies^16,17,22,25,36^. “Other studies”; Zhang et al., Martello et al., Bilan et al., and Bonfiglio et al. (I) Term enrichment analysis using Gene Ontology annotations, comparing proteins identified to be ADP-ribosylated to the total proteome. Relative score is derived from multiplication of logarithms derived from the enrichment ratio and the q-value. All displayed terms were significant with *q*<0.02, as determined through Fisher Exact Testing with Benjamini-Hochberg correction. “MF”; Molecular Functions, “CC”; Cellular Compartments, “BP”; Biological Processes. (J) As **I**, but using CORUM and InterPro annotations.

Overall, EThcD resulted in a higher number of identified ADP-ribosylated peptides, and EThcD was also more successful in localizing the modification to the correct amino acid (Figure 2A). Especially in case of the Lys-C samples, which naturally contained larger and thus harder-to-resolve peptides, EThcD outperformed pure ETD (Figure 2B). Upon investigation of all fragment ions contained within the MS/MS spectra, we did not observe a significant difference in average ion series coverage between ETD and EThcD (Figure S2E). However, EThcD demonstrated increased peptide bond coverage (Figure 2C), frequently covering >90% of all peptide bonds, and thus providing additional spectral evidence for true localization of ADP-ribosylation. We reason that the additional bond coverage is a direct result of the higher-collisional dissociation property of EThcD, which would be more likely to break specific bonds that would be resistant to pure ETD^33^.

When considering the contribution of each experiment to the total number of ADP-ribosylation peptides and sites identified (Figure 2D-E), we noted that the usage of trypsin and Lys-C resulted in highly complementary data. At the peptide level, 84.1% of all identifications were unique to either trypsin or Lys-C (Table S1), which remained 67.4% at the ADP-ribosylation site level (Table S2). Compared to trypsin and Lys-C, the overlap between ETD and EThcD was much larger, with EThcD providing most of the unique identifications.

In total, we identified 2,149 human proteins to be modified by ADP-ribosylation, with 57% modified on two or more sites (Figure 2F and Table S3). Ten or more ADP-ribosylation sites were mapped for 129 proteins, with the most sites identified on MKI67, harboring 91 sites in total. We identified 82% and 81% of modified sites and proteins reported in previous human ADP-ribosylation proteomics studies (Figure 2G-H, Tables S4 and S5), and additionally mapped 4,160 ADP-ribosylation sites and 782 ADP-ribosylation target proteins that have not been previously reported. Gene Ontology term enrichment analysis demonstrated that our expanded ADP-ribosylome corroborates the known functions of ADP-ribosylation (Table S6), with highly significant hits for nuclear localization, the DNA damage response, RNA metabolism, and chromatin remodeling (Figure 2I and S2F). Term enrichment analysis for protein complexes and structural domains showed frequent ADP-ribosylation of DNA binding proteins, helicases, the spliceosome, the ribosome, and histones (Figure 2J).

### Targeting preference of ADP-ribosylation

Because of the increased sequencing depth achieved through our extended analysis, we were able to profile the amino acid specificity of ADP-ribosylation to an unparalleled degree. The frequent modification of serine residues by ADP-ribosylation has recently been uncovered^22,36,38^, and we sought to validate whether the differential enzymatic digestions or fragmentation methods used here had any bearing upon these observations. Overall, the ADP-ribosylome profiled here largely corroborates the dominant targeting to serine residues (Figure 3A), with 88.7% of modifications occurring on serine residues, accounting for 97.2% of the measured ADP-ribosylation intensity (Table S2). Intriguingly, the three next-most abundantly modified amino acid residues were arginine, tyrosine, and histidine, with over a hundred sites detected and localized for each.

**Figure 3.**
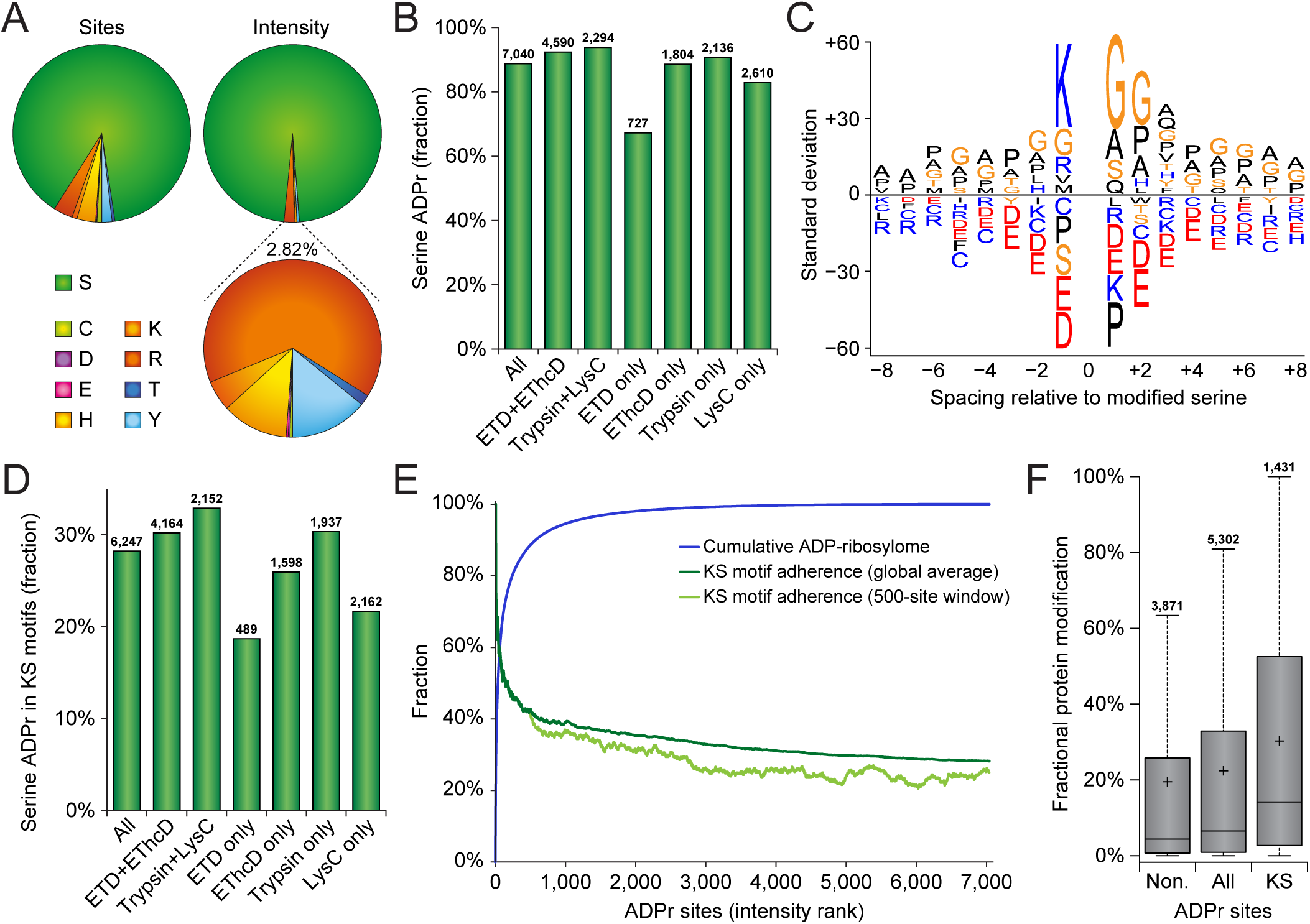
Targeting preference of ADP-ribosylation. (A) Pie-chart overview of the amino acid residue distribution of ADP-ribosylation sites (left pie-chart) and the distribution of ADP-ribosylation peptide peak intensity (top-right pie-chart). The bottom-right pie-chart represents a close-up of the 2.82% ADP-ribosylation intensity not occurring on serine residues. (B) Overview of the fraction of serine residues modified by ADP-ribosylation, in total and across several selections of ADP-ribosylation sites. Total numbers of ADP-ribosylation sites identified per selection are displayed above the bars. (C) IceLogo representation of the sequence context surrounding serine ADP-ribosylation sites. Amino acid residues displayed above the line were enriched, and those displayed below were depleted, compared to the sequence context of all serine residues derived from the same proteins. All displayed amino acids indicate significant changes as determined by two-tailed Student’s *t*-testing, *n*=6,245 serine ADP-ribosylation sites, *p*<0.05. (D) Overview of the fraction of serine ADP-ribosylation sites residing in KS motifs, in total and across several selections of ADP-ribosylation sites. Total numbers of serine ADP-ribosylation sites identified per selection are displayed above the bars. (E) Schematic depiction of the cumulative abundance of the ADP-ribosylome derived from ADP-ribosylation sites ranked by their peak intensities, with the total average and 500-site sliding window adherence to the KS motif plotted alongside. (F) Distribution of the fractional contribution of individual ADP-ribosylation sites to the total modification of the protein, for ADP-ribosylation sites not residing in KS motifs, all ADP-ribosylation sites, or ADP-ribosylation sites residing in KS motifs. Whiskers; 1.5× interquartile range (IQR), box limits; 3^rd^ and 1^st^ quartiles, center bar; median, + symbol; average. Numbers set above the top whisker correspond to the total number of serine ADP-ribosylation sites per grouping.

ADP-ribosylation sites identified through multiple modes of detection, i.e. with both fragmentation modes or with both enzymes, demonstrated an even higher adherence to serine residues (Figure 3B). We did not observe any large deviations in amino acid distribution for ADP-ribosylation sites detected exclusively in one experimental condition, except the relatively few ADP-ribosylation sites exclusively detected by ETD, where only 67.0% of modifications occurred on serine. Taken together, all methods investigated here highlight serine residues as the major ADP-ribosylation acceptor site.

As a second parameter for evaluation of technical performance, we assessed the sequence context surrounding the serine residues (Figure 3C). Primarily, lysine residues at the −1 position of the ADP-ribosylated serine residues, the so-called KS motifs, were highly prevalent in our data. Although glycine residues appear commonly nearby ADP-ribosylation sites, this can be attributed to their abundance in proteins^22^, unlike KS motifs. When considering the detection of KS motifs across the investigated experimental conditions, we observed 28.2% of all ADP-ribosylation sites to reside in KS motifs (Figure 3D), rising to 30.2% or 32.9% when considering those detected through both fragmentation modes or both enzymatic digestions, respectively. In terms of ADP-ribosylation sites exclusive to specific experimental conditions, those detected solely by ETD only adhered 18.7% to KS motifs, and those detected exclusively through LysC only adhered 21.7% to KS motifs.

Globally, when ranking ADP-ribosylation sites by their abundance, we found that the dynamic range of the ADP-ribosylome is very steep, with nearly two-third of ADP-ribosylation residing on the top 1% most-abundantly modified sites (Figure 3E and Table S2), and 92% of ADP-ribosylation residing on the top 10% most-modified sites. ADP-ribosylated serine residues in KS motifs frequently accounted for the most highly modified ADP-ribosylation sites (Figure 3E), with a steadily descending adherence across the one-third top-abundant sites. We analyzed the fractional contribution of individual ADP-ribosylation sites to the total modification of the respective proteins (Table S2), and used this information to investigate the intra-protein targeting of ADP-ribosylation to KS motifs (Figure 3F). Here, we observed that multiply-modified proteins that are ADP-ribosylated both on and outside of KS motifs, were significantly higher modified on KS motifs, suggesting a direct regulatory role for these motifs in attracting a high degree of ADP-ribosylation.

### Crosstalk between serine ADP-ribosylation and phosphorylation

With ADP-ribosylation and phosphorylation both intricately involved in regulation of cellular signaling, and able to modify a similar range of amino acid residues, we investigated our expansive repertoire of ADP-ribosylation sites for crosstalk between the two PTMs. Importantly, crosstalk can overall be divided into two categories (Figure 4A). First, co-targeting of the PTMs to the exact same residue, which is usually competitive but can also be sequential^39,40^. Second, co-modification of the protein by both PTMs simultaneously, which is usually cooperative and sequential.

**Figure 4.**
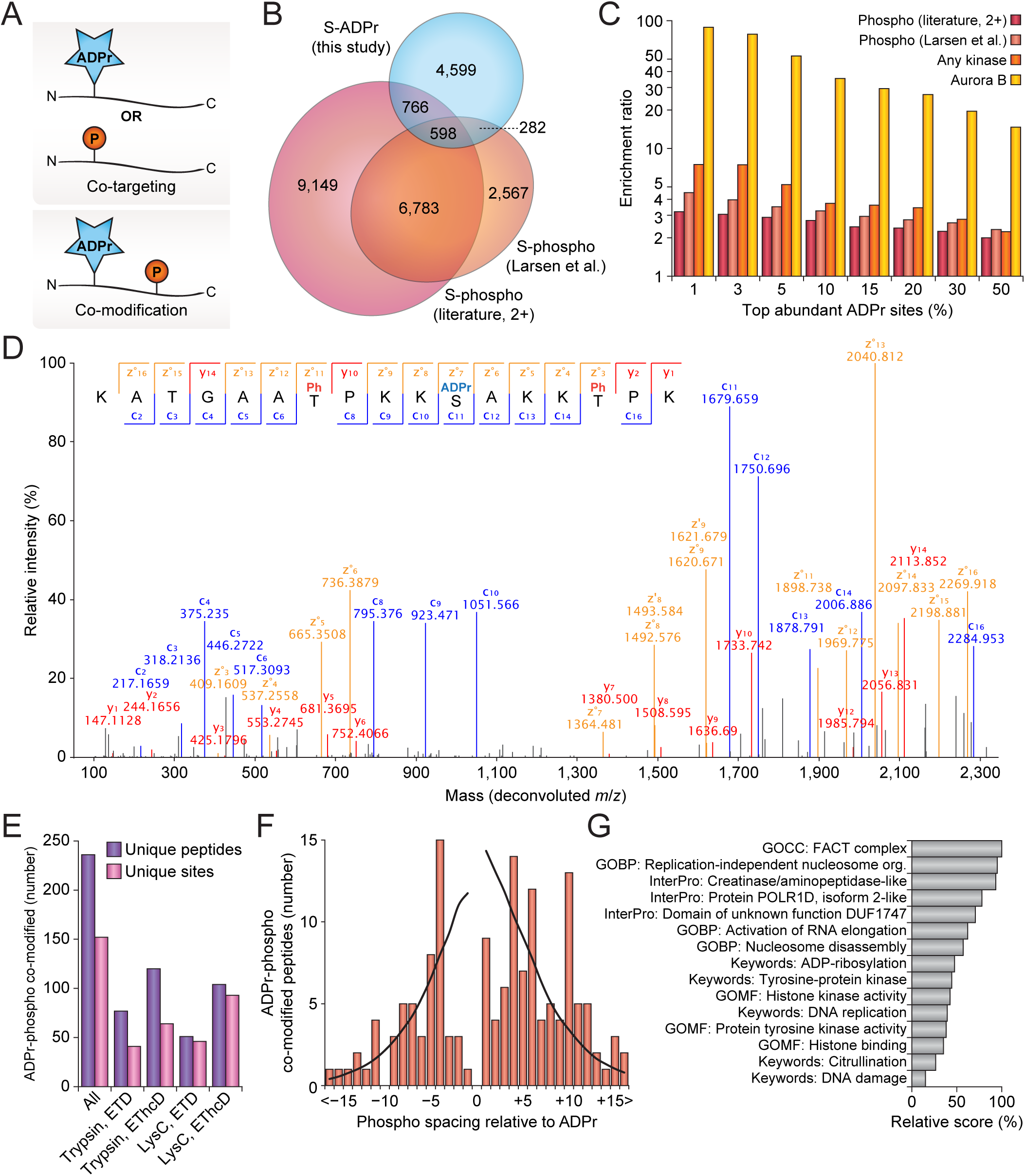
Crosstalk between ADP-ribosylation and phosphorylation. (A) Visualization of two distinct types of PTM crosstalk, either co-targeting or co-modification. (B) Scaled Venn diagram visualizing overlap between serine ADP-ribosylation sites identified in this study, and serine phosphorylation identified in a previous study^22^, or those identified by 2+ studies in the literature, as derived from PhosphoSitePlus (PSP)^41^. (C) PTM co-targeting enrichment analysis, visualizing overlap between top intensity serine ADP-ribosylation sites identified in this study with known serine phosphorylation sites, known kinase-regulated sites, and Aurora B regulated sites. All serine residues within the ADP-ribosylated proteins were used as a background, and enrichment ratios were determined through Fisher Exact Testing with Benjamini-Hochberg correction, *q*<0.02 in all cases. (D) Fully-annotated EThcD MS/MS spectrum, demonstrating confident localization of one ADP-ribosylation and two phosphorylation events. Blue; c-ions, red; y-ions, orange; z-ions, black; unassigned. (E) Overview of the number of ADP-ribosylation and phosphorylation co-modified peptides and sites identified in total, and across the four experimental conditions. (F) Visualization of the spacing of phosphorylation relative to ADP-ribosylation within co-modified peptides. The black line represents the fraction of instances where the phosphorylation spacing could theoretically be observed as derived from the distance to the termini for all ADP-ribosylated peptides identified in this study, and thus serves as a background profile. (G) Term enrichment analysis using various protein annotations, comparing ADP-ribosylation and phosphorylation co-modified sites to other ADP-ribosylation sites identified in this study. Relative score is derived from multiplication of logarithms derived from the enrichment ratio and the q-value. All displayed terms were significant with *q*<0.02, as determined through Fisher Exact Testing with Benjamini-Hochberg correction. “GO”; Gene Ontology, “GOMF”; GO Molecular Functions, “GOCC”; GO Cellular Compartments, “GOBP”; GO Biological Processes.

To investigate co-targeting of ADP-ribosylation and phosphorylation within our expanded dataset, we compared our ADP-ribosylation data to a previously published phosphorylation screen carried out in the same cell line and under similar conditions^22^, as well as global phosphorylation data^41^. ADP-ribosylation targeted to serine residues significantly overlapped to serine phosphorylation (Figure 4B and Table S6), with 26.4% of identified ADP-ribosylation sites also known to be targeted by phosphorylation. In line with what we previously described^22^, we found a significant overlap between H_2_O_2_-induced ADP-ribosylation and phosphorylation sites with known regulatory kinases (Figure 4C). Importantly, we found there was a significant correlation between the modification abundance of ADP-ribosylation sites and their propensity to overlap with phosphorylation, with the top-end ADP-ribosylation sites frequently overlapping with phosphorylation. This same trend was especially striking for ADP-ribosylation of serine residues usually modulated in their phosphorylation state through Aurora Kinase B, with a striking ~100-fold increased likelihood for the top 1% ADP-ribosylation sites to target these serine residues.

We next investigated co-modification of proteins by both ADP-ribosylation and phosphorylation at the same time. It is important to note that this requires peptides where both PTMs are simultaneously present and MS/MS spectra where both PTMs are confidently localized. The high resolution of our MS/MS spectra facilitated this analysis, as exemplified by identification and localization of one serine ADP-ribosylation and two phosphorylation events on a single Histone H1.4 peptide (Figure 4D). In total, 236 co-modified peptides were identified, corresponding to 152 unique combinations of ADP-ribosylation and phospho sites (Figure 4E and Table S7). Overall, similar to what we observed for ADP-ribosylated peptides in general, we found that EThcD was more successful at localizing both PTMs compared to pure ETD (Figure S3A), although all experimental conditions identified and localized dozens of co-modified peptides. In terms of proteolytic enzymes, we found that co-modified peptides generated by Lys-C digestion were more frequently identified and less likely to lead to identification of redundant peptides (Figure 4E), which follows the larger peptide sequences generated upon LysC digestion. Compared to the ADP-ribosylated residue, phosphorylation was frequently observed to occur at a spacing of −4, +4, +6, or +10 residues (Figure 4F), although there was no strong enrichment in any specific position. Rather, the even-numbered spacing could hint at these residues being solvent-exposed considering that ADP-ribosylation and phosphorylation both target disordered regions^22,42^. We observed numerous peptides derived from histones to be co-modified (Table S7), suggesting that histones could be a preferred target of co-regulation by ADP-ribosylation and phosphorylation crosstalk. Term enrichment analysis of proteins found to be preferentially co-modified by ADP-ribosylation and phosphorylation highlight transcriptional elongation, chromatin remodeling, DNA replication, the DNA damage response, tyrosine kinase activity, and ADP-ribosylation itself (Figure 4G and Table S6).

### ADP-ribosylation on non-serine residues

Although the majority of our data entailed ADP-ribosylation of serine residues, our diversified experimental approach allowed us to additionally localize and profile an unprecedented number of ADP-ribosylation events on other amino acid residues. Next to serine, the three most abundantly modified amino acid residues were histidine, arginine, and tyrosine, in both number of modified residues and the overall fraction of the ADP-ribosylome residing on those residues. In total, 705 sites were identified and localized to these three residue types, encompassing a combined 2.57% of the ADP-ribosylome (Figure 5A and Table S2).

**Figure 5.**
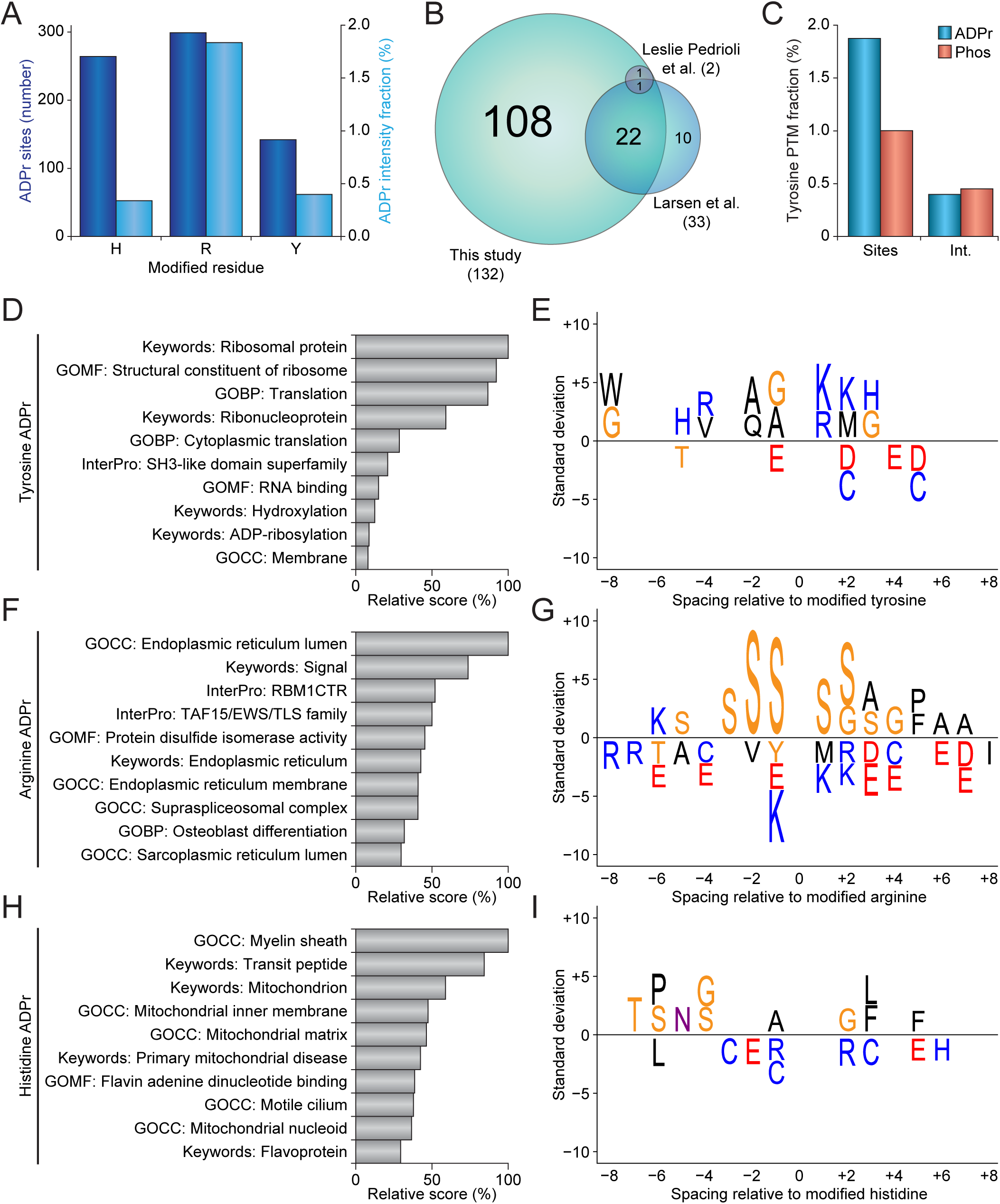
ADP-ribosylation on non-serine residues. (A) Overview of the number of ADP-ribosylation sites modifying histidine, arginine, and tyrosine residues (dark blue bars), and the fraction of total ADP-ribosylation intensity residing on histidine, arginine, and tyrosine residues (light blue bars). (B) Scaled Venn diagram visualizing overlap between tyrosine ADP-ribosylation sites identified in this study and in two other proteomics studies reporting tyrosine ADP-ribosylation^22,43^. Only tyrosine ADP-ribosylation sites confidently localized using ETD-type fragmentation were considered. (C) Comparison of the numerical and intensity fractions of ADP-ribosylation and phosphorylation identified on tyrosine residues. (D) Term enrichment analysis using various protein annotations, comparing tyrosine ADP-ribosylation sites to all other ADP-ribosylation sites identified in this study. Relative score is derived from multiplication of logarithms derived from the enrichment ratio and the q-value. All displayed terms were significant with *q*<0.02, as determined through Fisher Exact Testing with Benjamini-Hochberg correction. “GO”; Gene Ontology, “GOMF”; GO Molecular Functions, “GOCC”; GO Cellular Compartments, “GOBP”; GO Biological Processes. (E) IceLogo representation of the sequence context surrounding tyrosine ADP-ribosylation sites. Amino acid residues displayed above the line were enriched, and those displayed below were depleted, compared to the sequence context of all tyrosine residues derived from the same proteins. All displayed amino acids indicate significant changes as determined by two-tailed Student’s *t*-testing, *n*=142 tyrosine, *n*=299 arginine, and *n*=264 histidine ADP-ribosylation sites, *p*<0.05. (F) As **D**, but for arginine ADP-ribosylation. (G) As **E**, but for arginine ADP-ribosylation. (H) As **D**, but for histidine ADP-ribosylation. (I) As **E**, but for histidine ADP-ribosylation.

Since tyrosine ADP-ribosylation has been previously reported^43,44^, we first performed a comparison of our tyrosine ADP-ribosylation sites to other known sites that were detected through unbiased proteomics approaches relying on ETD-based fragmentation (Figure 5B). We identified 71% of previously mapped tyrosine ADP-ribosylation sites, and further expanded the number of tyrosine ADP-ribosylation sites by >300%, showing that our experimental approach especially facilitates identification of ADP-ribosylation sites with low-abundance. Intriguingly, we observed a tendency for both the number of tyrosine ADP-ribosylation sites and the overall intensity of the ADP-ribosylome to be very comparable to that of tyrosine phosphorylation when profiled at similar depth, in the same cell line, and in response to the same cellular treatment (Figure 5C).

With tyrosine also a well-known target of phosphorylation, we performed a co-targeting crosstalk analysis similar to serine ADP-ribosylation (vide supra). A high degree of overlap was observed compared to previously reported tyrosine phosphorylation sites (Figure S3B and Table S6), with 42% of tyrosine ADP-ribosylation sites known to also be targeted by phosphorylation. A significant overlap was observed with tyrosine ADP-ribosylation sites reported recently^22,43^, and the overlap between tyrosine ADP-ribosylation mapped in this study and tyrosine phosphorylation was also found to be significant (Figure S3C). The overlap between tyrosine ADP-ribosylation and phosphorylation was of a similar magnitude as in the case of serine modification, thus hinting at a similar regulatory crosstalk at the tyrosine level.

We next investigated whether we could derive new insight from the residue-specific ADP-ribosylation in regards to biological function. To this end, we used data generated in this study, and to increase statistical accuracy we included tyrosine, arginine, and histidine ADP-ribosylation data from our previous study^22^. For tyrosine ADP-ribosylation, we mainly noted a high propensity for these events to modify ribosomal proteins (Figure 5D and Table S6), with no strong enrichment for any other processes in comparison to general ADP-ribosylation. In terms of sequence context, no strong enrichment was observed (Figure 5E), with the most significant motif suggesting a lysine residue at the +1 position of the modified tyrosine; i.e. a YK motif. For arginine ADP-ribosylation, enriched processes were mainly related to the endoplasmic and sarcoplasmic reticulum, along with modification of RNA binding domains (Figure 5F and Table S6). The protein sequence context surrounding ADP-ribosylated arginines primarily suggested a strong enrichment for serine residues at all flanking positions (Figure 5G). For histidine ADP-ribosylation, the enriched terms suggested enrichment in the mitochondrion, myelin sheath, and flavoproteins (Figure 5H and Table S6), while the sequence context did not reveal any striking enrichments (Figure 5I).

Collectively, our residue-specific analysis suggested that ADP-ribosylation may be targeted differentially depending on subcellular localization, or in relation to specific protein complexes. To further validate and visualize our observations, we subjected all proteins to functional clustering analysis through the STRING database, and then clustered interconnected proteins by the major function we derived for ADP-ribosylation of the specific amino acid types (Figure S4). Overall, 88.6% of the 481 proteins clustered together, and although all of these were modified on at least one histidine, arginine, or tyrosine, the majority of ADP-ribosylation still occurred on serine residues for proteins not specifically associated with non-serine ADP-ribosylation clusters through our previous analysis (Figure 5D-E). However, a modest enrichment could be observed for tyrosine residues in the ribosome cluster, and strong enrichments for modification of histidine residues in the mitochondrion cluster, and arginine residues in the endoplasmic reticulum cluster, hinting at specific biological roles and regulations for tyrosine, arginine, and histidine ADP-ribosylation events.

## DISCUSSION

In this study, we optimized our MS-based proteomics methodology for the identification and localization of ADP-ribosylation sites. To this end, we expanded upon the established Af1521 workflow by incorporating separate trypsin and Lys-C digestions, optimizing several stages of the purification, and using both ETD and EThcD fragmentation of precursors. Overall, this allowed us to identify >7,000 unique ADP-ribosylation sites modifying >2,000 proteins, corresponding to over one-third of the nuclear proteome.

Using exclusively Lys-C to digest proteins generates larger, more hydrophobic, and higher charged peptides, because contrary to tryptic digestion there is no cleavage C-terminal of arginine residues. We expected the higher charge state to aid in identification of the peptides, as we previously observed that ADP-ribosylated precursors with a charge state of 2 are very difficult to resolve with ETD-type fragmentation^22^. Indeed, we found that Lys-C digestion resulted on average in somewhat larger ADP-ribosylated peptides, higher charged precursors, and higher spectral assignment scores and lower posterior error probabilities. However, owing to the increased size of the peptides, high-confidence localization of ADP-ribosylation proved more challenging (Figures 2B and S4C), ultimately leading to a virtually identical identification rate of the trypsin- or Lys-C-derived peptides.

Nonetheless, we found that the tryptic and Lys-C digests resulted in the generation of highly complementary peptides, with many of them harboring ADP-ribosylation events that were exclusive to either one or the other, and over two-thirds of all ADP-ribosylation sites uniquely identified through one enzymatic digest. Usage of multiple digestion enzymes has previously been reported to increase protein sequence coverage^29,37,45^, with certain digestion strategies precluding regions that would be represented by peptides that are either too short or too long for mass spectrometric analysis, e.g. arginine-rich protein regions when using tryptic digestion. Contrary to total protein analysis, PTM analysis is restricted to finding specific and often low-abundant peptides, and we reason that because of this the added effect of using different digestion strategies is magnified. However, usage of multiple proteases comes with an increased demand of mass spectrometric analyses. Thus, usage of either trypsin or Lys-C digestion for analysis of ADP-ribosylated peptides could be considered as another dimension of sample fractionation.

Previously, we relied on pure ETD to identify and localize all ADP-ribosylation events, after establishing that HCD fragmentation usually prohibits reliable localization of ADP-ribose when using unrestricted database searches^22^. Since EThcD is in essence a combination of ETD and HCD, we initially worried that the inclusion of the HCD dimension could partially nullify the non-ergodic propensity of ETD fragmentation. Others have demonstrated application of EThcD for detection of ADP-ribosylated peptides^36^, but used relatively high levels of supplemental activation (SA) energy. In our initial optimization, we used relatively low amounts of SA energy in order to establish a good balance between ETD- and HCD-driven fragmentation. Indeed, we observed that at the lowest SA energy, the fragmentation behavior of ADP-ribosylated peptides essentially mimicked pure ETD fragmentation. Even towards the highest SA energy we used, we saw increasingly high localization probability of the ADP-ribose, and importantly the ADP-ribose remained attached to the peptides even at the higher SA energies. This suggests that the peptide fragmentation remains primarily driven by ETD, with the HCD dimension assisting in breaking peptide bonds that are resistant to ETD. Nonetheless, we observed a drop in identification efficiency at the highest level of SA energy we tested, which could be related to an on average decreasing spectral quality as SA energy was increased, likely a result of increasing spectral complexity with larger fractions of the peaks remaining unassigned.

Overall, we found that using EThcD fragmentation with a low amount of SA energy increased both identification and localization of ADP-ribosylation, compared to pure ETD fragmentation. However, compared to using either Lys-C or trypsin to digest the samples, we did not observe a lot of complementarity between analyzing samples with either ETD or EThcD, with EThcD generally outperforming ETD. Thus, whereas analyzing half of the samples with trypsin or Lys-C digestions could yield a higher number of identifications rather than analyzing full samples with either one of the enzymes alone, we surmise that it would in most cases be optimal to analyze all samples exclusively with EThcD.

Although we initially expected that EThcD would aid in dissociation of charge-reduced ADP-ribosylated peptide precursors, we did not observe this to a notable degree. Indeed, it has previously been reported that EThcD is not necessarily optimal for this purpose^46^. A potential optimization of our method could involve application of activated ion ETD (AI-ETD), which is based on infrared photoactivation^46^, and is very efficient at dissociation both precursors and charge-reduced precursors. AI-ETD has been used for total proteome analysis as well as for phosphoproteomics, and has proven superior to not only ETD alone, but also to EThcD^46,47^, indicating that this could be a valuable technology for studying ADP-ribosylation.

With the increased depth of sequencing entailed within our dataset, along with the overall higher level of spectral quality, we were able to more confidently derive biological phenomena. We could readily confirm all known system-wide properties of ADP-ribosylation, including a predominantly nuclear localization, modification of a plethora of proteins involved in DNA repair and chromatin remodeling, preferential modification of serine residues, targeting of serine residues residing in KS motifs, and a significant overlap with phosphoserines. Moreover, we were able to statistically verify new phenomena, including co-targeting of tyrosine residues by ADP-ribosylation and phosphorylation, simultaneous co-modification of proteins by ADP-ribosylation and phosphorylation, and functional associations with ADP-ribosylation of histidine, arginine, and tyrosine residues.

Crosstalk between phosphorylation and ADP-ribosylation has been previously described^44,48^, and we could observe a notable degree of this within our data. Importantly, we found that there is an abundance-driven co-targeting for serine residues, with the most abundant ADP-ribosylation events having the highest degree of overlap with phosphoserines, and specifically those for which regulatory kinases are known. Biologically, this suggests a regulatory role for ADP-ribosylation with the targeting of these residues, as the prompt modification of these serine residues would prohibit phosphorylation from modifying them. Furthermore, we observed a similar effect for tyrosine ADP-ribosylation, which significantly overlapped with previously reported tyrosine phosphorylation events. The observed degree of tyrosine co-targeting was comparable to serine co-targeting, which could similarly suggest a regulatory role for ADP-ribosylation in precluding tyrosine phosphorylation.

Identification of peptide co-modification is innately technically challenging, because peptides have to be resolved that harbor both PTMs, both PTMs have to be localized, and the analysis is by nature subject to the potentially limited abundance of both PTMs. Nonetheless, we were able to identify 236 ADP-ribose and phosphorylation co-modified peptides, corresponding to 152 unique sites, and accounting for ~1% of the total ADP-ribosylome. Systems-wide mapping of these co-modified peptides suggests that, other than co-targeting of the same residues, ADP-ribosylation and phosphorylation may frequently be targeted to the same regions of proteins. Although we did not observe a predominant spacing or motif between the two PTMs, their spatiotemporal co-existence and preferential targeting to disordered regions of proteins could suggest another layer of regulatory crosstalk between ADP-ribosylation and phosphorylation.

When considering ADP-ribosylation targeted to amino acid residues other than serine, we observed frequent modification of histidine, arginine, and tyrosine residues. Modification of all three of these amino acid residues by ADP-ribosylation has been previously reported^11,15,43^, although only modification of arginine residues has been more widely studied. We carried out our experiments in HeLa cells exposed to hydrogen peroxide, which does not necessarily equate a model system specifically tuned for detection of specific types of amino acid residues being ADP-ribosylated. Nonetheless, the unprecedented depth of our dataset allowed us to observe uniquely enriched biological pathways for each type of amino acid residue. For tyrosine residues, we observed a propensity for modification of ribosomal proteins. It should be mentioned that overall tyrosine ADP-ribosylation occurred on high-abundant proteins, and the modified proteins were frequently also modified on other types of amino acid residues. For arginine residues, preferential modification of proteins associated with the endoplasmic reticulum was observed, which is consistent with the reported localization of human ARTC1 to the endoplasmic reticulum^49^. Sequence context analysis highlighted a relatively high degree of serine residues flanking the modified arginine, potentially because of a propensity for arginine ADP-ribosylation to target RNA-binding proteins rich in serine and arginine residues. For histidine residues, modification was found on proteins associated with the mitochondrion. Histidine phosphorylation has previously been observed to modify mitochondrial proteins, and has been linked to regulation of mitochondrial function^50^. Analogous to histidine phosphorylation, ADP-ribosylation of histidines could potentially be involved in mitochondrial regulation.

Collectively, we expand knowledge regarding the human ADP-ribosylome using an augmented and unbiased enrichment strategy based on the Af1521 macrodomain. We find that EThcD outperforms pure ETD fragmentation for identification and localization of ADP-ribosylation, whereas digestion using multiple proteases identifies complementary sites. Taken together, we observe modification of over one-third of the nuclear proteome by ADP-ribosylation in response to DNA damage, and our comprehensive dataset of ADP-ribosylation sites will aid in understanding the system-wide scope of this pivotal post-translational modification.

## AUTHOR CONTRIBUTIONS

S.C.L and I.A.H. prepared all samples, performed all experiments, and measured all samples on MS. I.A.H., S.C.L., and M.L.N. processed all MS raw data. I.A.H. and S.C.L. optimized MS methodology and performed all bioinformatics and statistical analyses. M.L.N conceived and supervised the project. I.A.H., S.C.L., and M.L.N. wrote the manuscript.

## ACKNOWLEDGEMENTS

The work carried out in this study was in part supported by the Novo Nordisk Foundation Center for Protein Research, the Novo Nordisk Foundation (grant agreement numbers NNF14CC0001 and NNF13OC0006477), The Danish Council of Independent Research (grant agreement numbers 4002-00051, 4183-00322A and 8020-00220B), and The Danish Cancer Society (grant agreement R146-A9159-16-S2). I.A.H. is supported by the European Molecular Biology Organization (grant agreement number ALTF 503-2016). We would like to acknowledge the lab of Michael O. Hottiger for the expression and purification of recombinant human PARG (University of Zurich), and we thank members of the NNF-CPR Mass Spectrometry Platform for instrument support and technical assistance.

## SUPPLEMENTARY FIGURE LEGENDS

**Figure S1.**
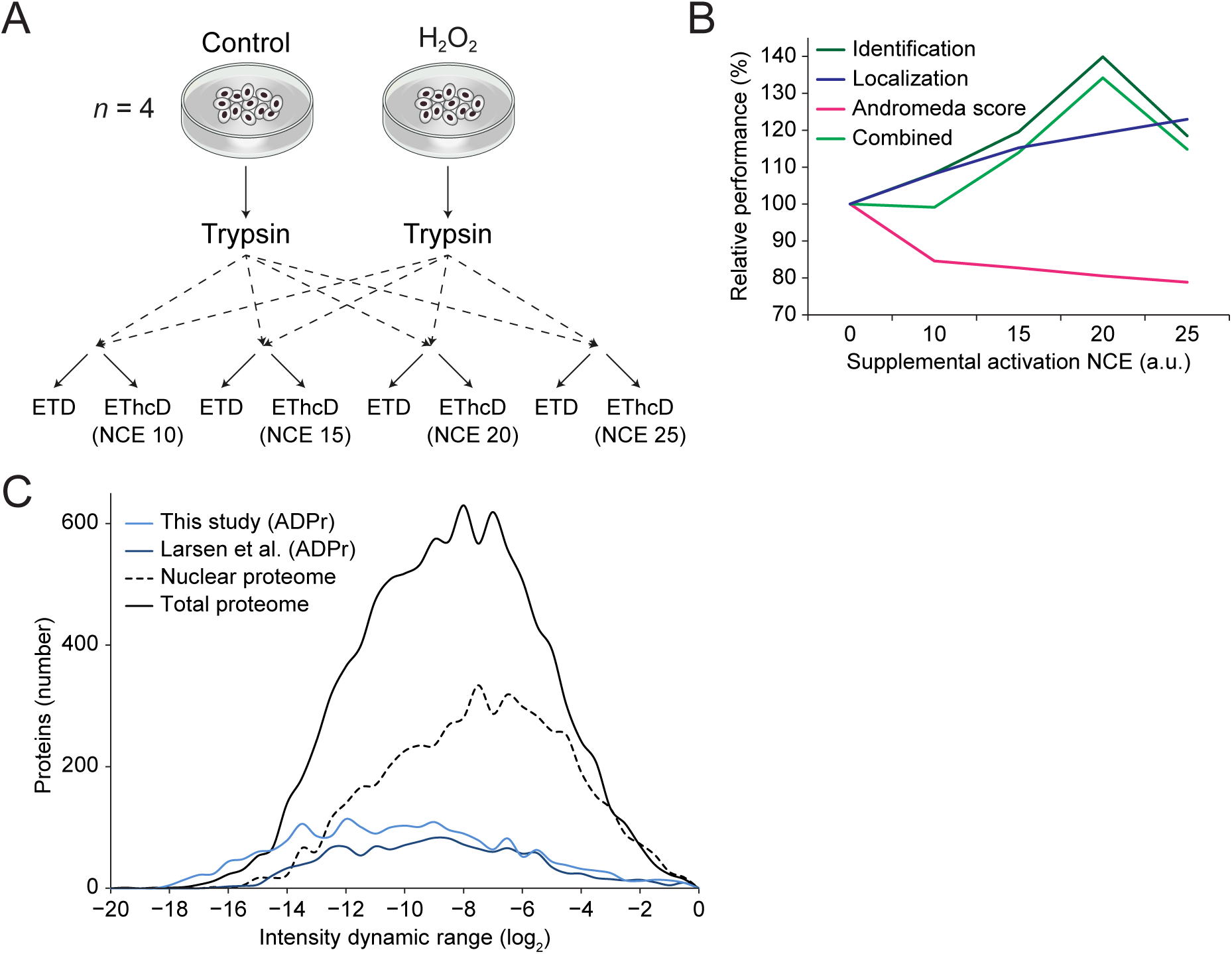
(A) Experimental design for initial experiments performed to evaluate different levels of EThcD supplementary activation (SA) energy. Cell cultures were performed in quadruplicate; *n*=4, and all cells were either left untreated or treated with H_2_O_2_ to induce ADP-ribosylation. All samples were prepared through the trypsin workflow, and final samples were split in halves and measured back-to-back as outlined. (B) Schematic overview of various attributes as profiled across the different SA energies. Values represent the number of total ADP-ribosylated peptide identifications, the average ADP-ribose localization probabilities, and the average Andromeda scores, in each instance normalized between the EThcD half of the sample and the ETD half of the sample. The combined score was derived by multiplication of the three other values. (C) Dynamic depth of sequencing graph depicting the range of protein intensity values determined in this study, plotted against the number of proteins identified, and compared to various other studies^22,37^. Protein intensity values were derived from all peak intensities of all corresponding peptides. All studies were normalized to have the most-abundant proteins align at the right side of the plot.

**Figure S2.**
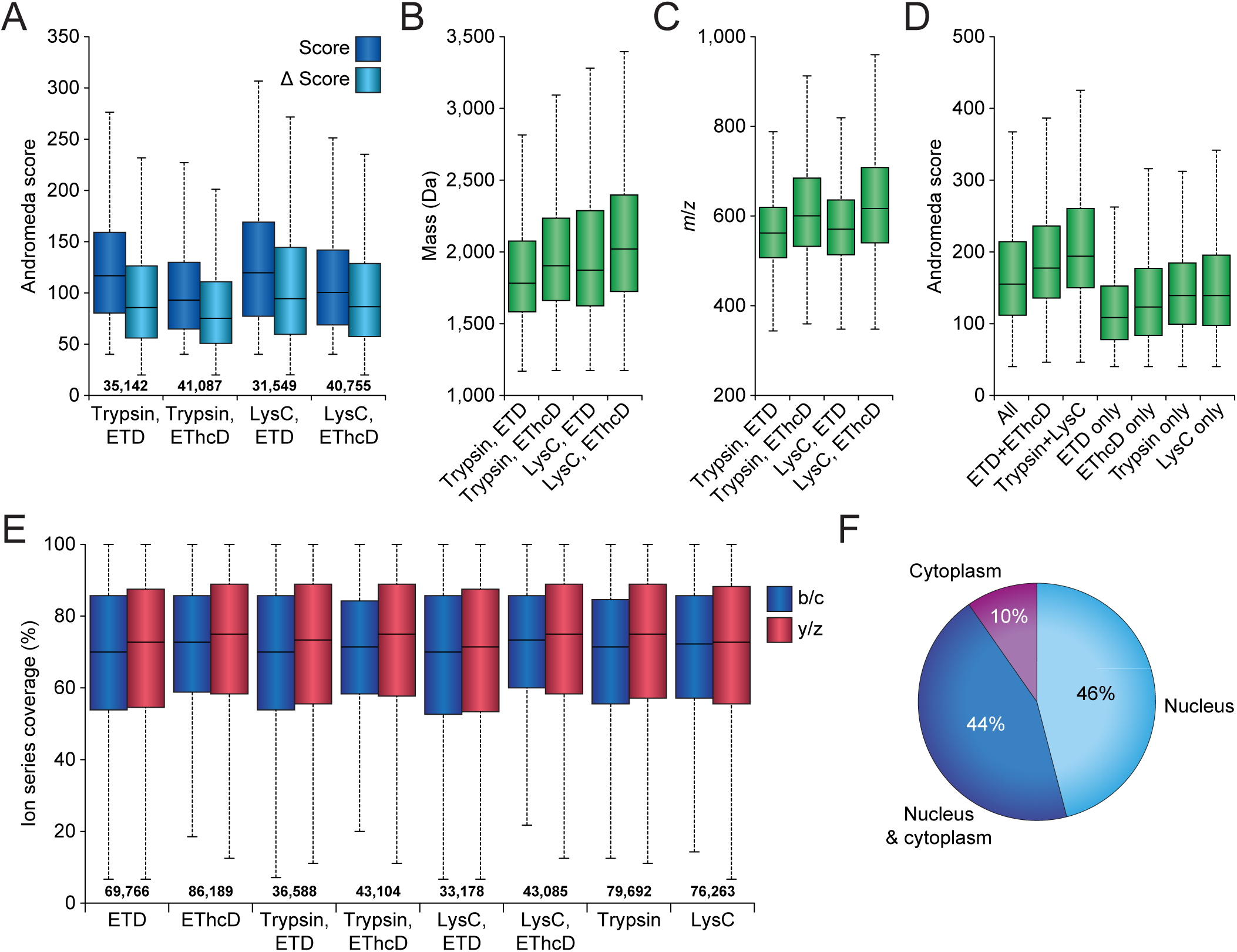
(A) Boxplot graph visualizing distribution of the Andromeda score and delta score for ADP-ribosylation PSMs, as measured across the four experimental conditions. Whiskers; 1.5× interquartile range (IQR), box limits; 3^rd^ and 1^st^ quartiles, center bar; median. Numbers set above the horizontal axis correspond to the number of identified ADP-ribosylation PSMs. (B) As **A**, but for peptide mass. (C) As **A**, but for precursor *m/z*. (D) As **A**, but for ADP-ribosylation sites, and additionally for subsets of sites identified through multiple experimental conditions, or exclusive to certain experimental conditions. (E) Ion series coverage analysis, displaying the fraction of the maximum observable ion series for which peaks were detected and assigned, divided between C-terminal (b/c) and N-terminal (y/z) fragment series. Whiskers; 1.5× interquartile range (IQR), box limits; 3^rd^ and 1^st^ quartiles, center bar; median. Numbers set above the horizontal axis correspond to the number of identified ADP-ribosylation PSMs. (F) Pie-chart visualization of the subcellular localization of proteins identified to be ADP-ribosylated.

**Figure S3.**
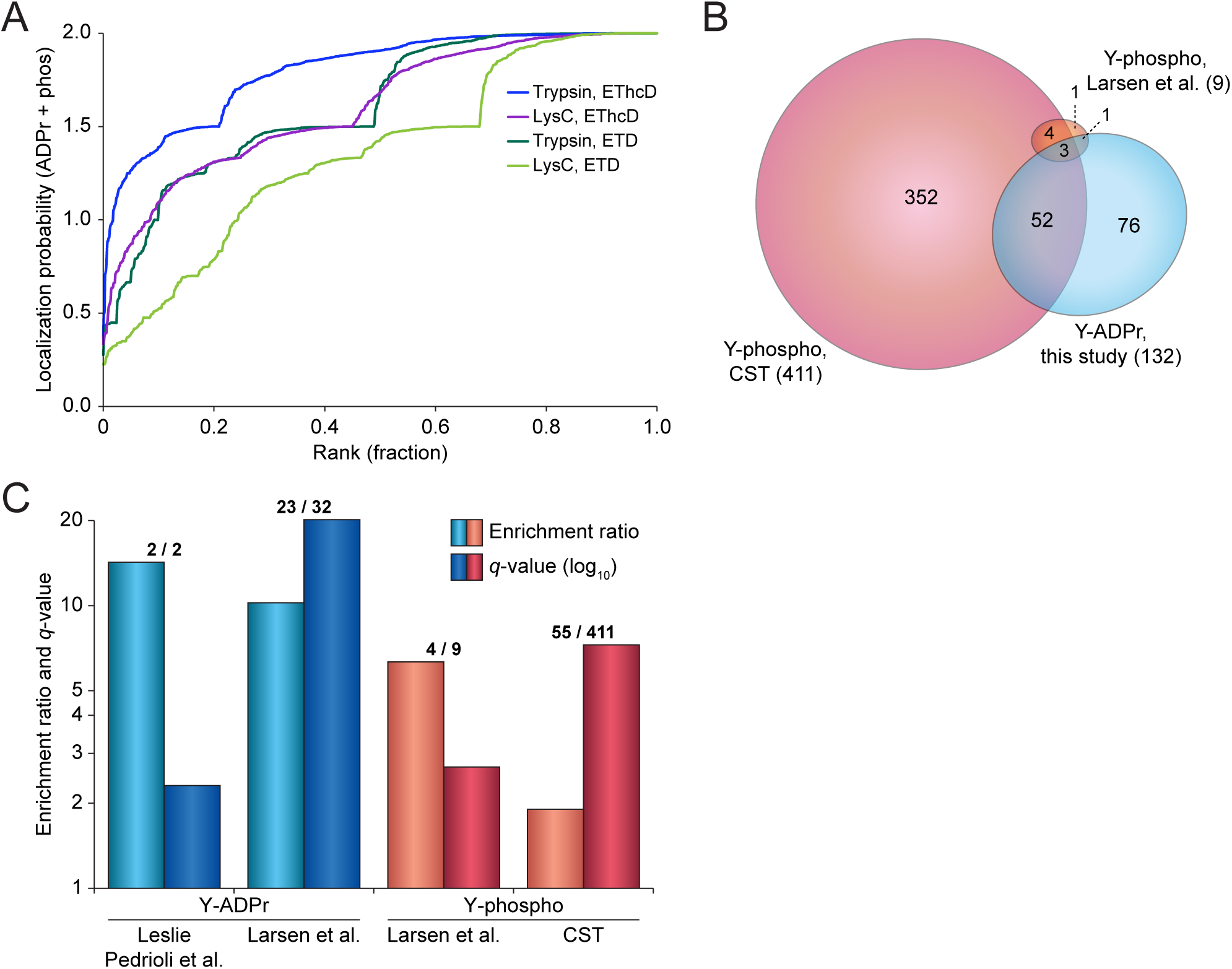
(A) Combined ADP-ribosylation and phospho localization probability plotted against the ranked fraction of all peptide-spectrum-matches (PSMs) resulting from the four different approaches. Individual probabilities for ADP-ribosylation and phospho were summed, and co-modified peptides and sites were only derived when both PTMs were individually localized at over 0.9 probability. (B) Scaled Venn diagram visualizing overlap between tyrosine ADP-ribosylation sites identified in this study, and tyrosine phosphorylation identified in one previous study^22^, or those identified and reported by Cell Signaling Technology (CST)^41^. (C) PTM co-targeting enrichment analysis, depicting overlap and significance of overlap between tyrosine ADP-ribosylation sites identified in this study, tyrosine ADP-ribosylation sites identified in two other studies^22,43^, and tyrosine phosphorylation identified in one previous study^22^ or by Cell Signaling Technology^41^. All tyrosine residues within the ADP-ribosylated proteins were used as a background, and enrichment ratios and *q*-values were determined through Fisher Exact Testing with Benjamini-Hochberg correction. Numbers set above the bars correspond to the intersection between tyrosine ADP-ribosylation identified in this study, in relation to the total number of tyrosine PTMs reported in the other studies.

**Figure S4.**
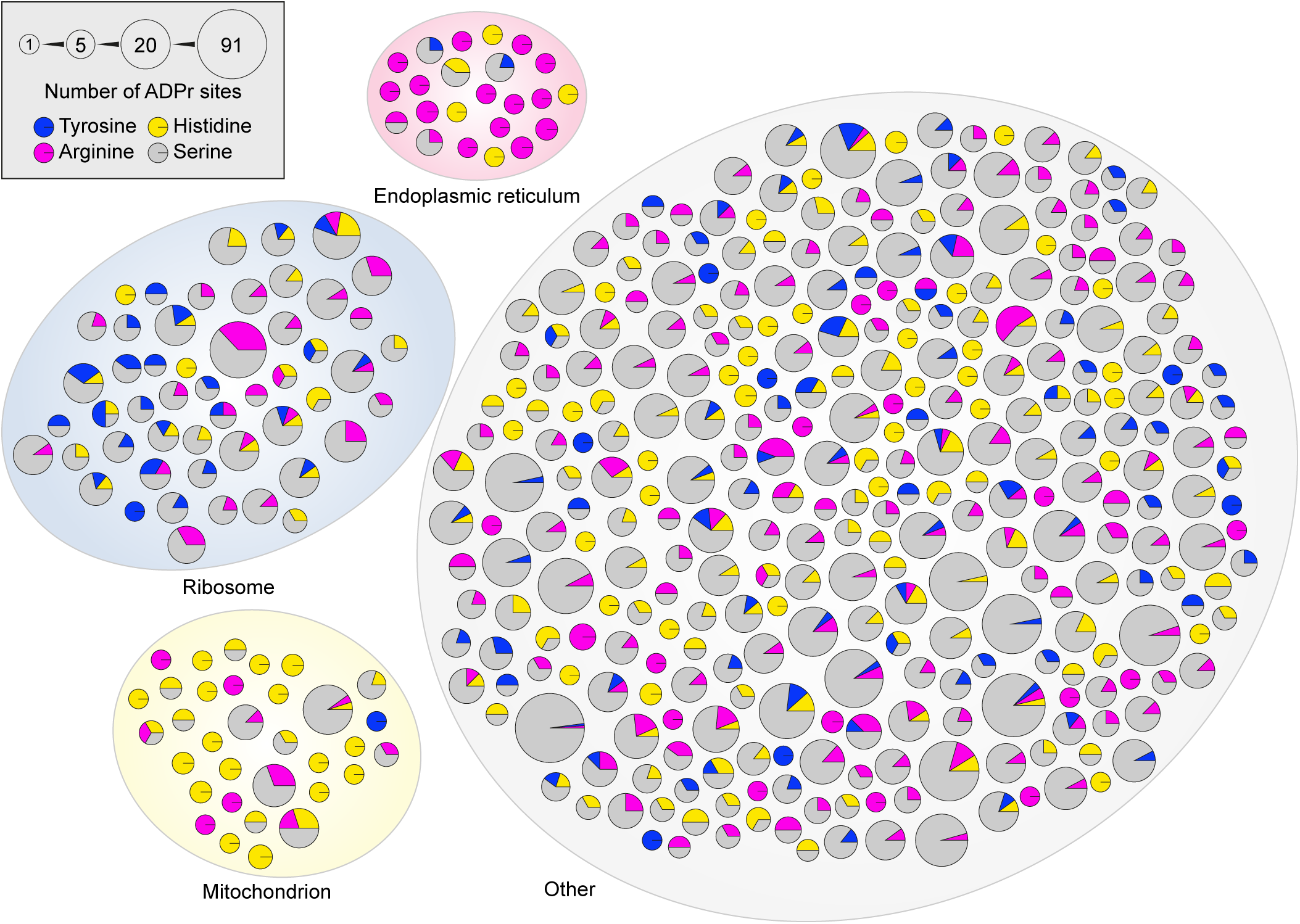
STRING network analysis of all ADP-ribosylated proteins modified on at least one histidine, arginine, or tyrosine residue. Default STRING clustering settings were used (clustering confidence >0.4), proteins not clustered were omitted from the network, and edges were hidden. Proteins were curated based on the number of ADP-ribosylation sites and the type of residues modified by ADP-ribosylation, as indicated. ADP-ribosylation sites on amino acid residues other than serine, histidine, arginine, and tyrosine, were omitted. Proteins annotated as part of the mitochondrion, ribosome, or endoplasmic reticulum, were clustered and moved out of the main network to assist visualization.

## METHODS

### Cell culture

HeLa cells (CCL-2) were acquired from the American Type Culture Collection, and cultured in Dulbecco’s modified Eagle’s medium (Invitrogen) supplemented with 10% fetal bovine serum and penicillin/streptomycin (100 U/mL; Gibco) at 37°C and 5% CO_2_. Cells were not routinely authenticated. Cells were routinely tested for mycoplasma contamination. Per replicate, 100 million HeLa cells were cultured for the initial EThcD energy optimization experiments, and 300 million HeLa cells were cultured for the main experiments. To induce ADP-ribosylation in HeLa, cells were treated with 1 mM H_2_O_2_ (Sigma Aldrich) for 10 min in PBS at 37°C.

### Cell lysis

To maximize protein recovery and minimize loss of labile PTMs, we used a lysis procedure reliant on the chaotropic salt guanidine-HCl. After H_2_O_2_ treatment, cells were washed twice with ice-cold PBS, and collected by gentle scraping at 4°C. Cells were pelleted in a swing-out centrifuge with delayed deceleration at 4°C, for 3 min at 500*g*. PBS was decanted, and the cell pellets were directly lysed in 10 pellet volumes of lysis buffer (6 M guanidine-HCl, 50 mM TRIS, pH 8.5). Rapid lysis was achieved by alternating vigorous vortexing and vigorous shaking of the samples, for 5 seconds per cycle, and 30 seconds in total, after which lysates were snap frozen using liquid nitrogen. Frozen lysates were stored at −80°C until further processing.

### Protein digestion

Cell lysates were thawed at room temperature, and homogenized using sonication with three 15 second pulses at 30 W. Homogenized lysates were supplemented with 5 mM TCEP and 5 mM chloroacetamide (CAA), and incubated for 1 hour at 30°C. After reduction and alkylation, samples were split in two equal halves, and digested using Lysyl Endopeptidase (Lys-C, 1:100 w/w; Wako Chemicals) for 3 hours, at room temperature and without sample agitation. After initial digestion, samples were diluted with three volumes of ice-cold 50 mM ammonium bicarbonate (ABC), after which they were allowed to slowly warm up to room temperature over a period of 30 min. Subsequently, half of the samples were further digested using modified sequencing grade Trypsin (1:200 w/w; Sigma Aldrich), and the other half was re-digested with Lys-C, in both cases overnight at room temperature and without sample agitation. Post-digestion, samples were acidified using trifluoroacetic acid (TFA) to a final concentration of 0.5% (v/v). Samples were cleared from precipitates by centrifugation in a swing-out centrifuge at 4°C, for 30 min at 4,250*g*. Peptides were purified by loading samples onto reversed-phase C18 cartridges (SepPak Classic, 360 mg sorbent, Waters), which were pre-activated with 5 mL ACN, and equilibrated 3× with 5 mL of 0.1% TFA. Sample loading was accelerated using a vacuum manifold, utilizing a low vacuum of 2/3^rd^ atmospheric pressure. After loading, cartridges were washed with 3× with 5 mL of 0.1% TFA, after which peptides were eluted using 4 mL of 30% or 40% ACN in 0.1% TFA, for peptides digested with either trypsin or Lys-C, respectively. Eluted peptides were frozen at −80°C overnight, in 50 mL tubes with small holes punctured in the caps, after which they were dried to completion by lyophilization over the course of 96 h. Dried peptides were stored at −20°C until further processing.

### Purification of Af1521 macrodomain

To purify ADP-ribosylated peptides, we utilized a GST-tagged Af1521 macrodomain, produced in-house essentially as described previously^22^. Competent BL21(DE3) were incubated with GST-Af1521 plasmid for 30 min, after which they were heat shocked at 42°C for 45 s, incubated on ice for 2 min, allowed to recover for 1 hour, plated on ampicillin LB plates, and incubated overnight at 37 °C. Four colonies were picked and transferred into LB media, growing overnight at 37°C while shaking. The starter culture was diluted 1:20 and grown to a 600-nm optical density of 0.60, after which protein expression was induced through addition of IPTG to a final concentration of 0.5 mM for 6 h at 30°C. Bacteria were collected by centrifugation, washed twice with PBS, and ultimately 100 mL worth of bacteria-containing LB was centrifuged per 50 mL tube. Bacterial pellets were frozen at −80 °C until further processing. Bacteria were lysed using BugBuster^®^ Protein Extraction Reagent (Merck) according to the manufacturer’s instructions. GST-tagged Af1521 was purified from bacterial lysates using glutathione Sepharose 4B beads (Sigma Aldrich). 2.5 mL of beads (dry volume) were incubated with the bacterial lysate for 4 h at 4°C in a head-over-tail mixer, after which they were washed five times using PBS. Tubes were changed after the 2^nd^ and 4^th^ wash steps to minimize contaminant carryover, and pelleting of beads was performed in a swing-out centrifuge with delayed deceleration to minimize loss of beads, at 4°C for 3 min at 500*g*. After the final wash, 1 mL of beads were stored per 15 mL tube, with PBS entirely filling up the tube, and supplemented with 10 mM sodium azide to retard microbial growth. Beads were stored at 4°C until use, for up to three months.

### Reduction of ADP-ribose polymers

Lyophilized peptides were dissolved in IP buffer (50 mM TRIS pH 8.0, 1 mM MgCl_2_, 250 μΜ DTT, and 50 mM NaCl) for two replicates, and in 5× IP buffer (250 mM TRIS pH 8.0, 5 mM MgCl_2_, 1.25 mM DTT, and 250 mM NaCl) for the other two replicates, in order to assist dissolution of potentially more hydrophobic peptides. However, after investigation of the samples no significant differences were observed between dissolution of peptides in 1× and 5× IP buffer, and therefore the two sets of two replicates were subsequently treated as one set of four replicates. As not all peptides dissolved at alkaline pH after being purified at acidic pH, samples were cleared by centrifugation in a swing-out centrifuge at room temperature, for 30 min at 4,250*g*. The concentration of the peptides was determined using UV spectroscopy, and was approximately 1/4^th^ of the initial total protein amount. ADP-ribose polymers were reduced to monomers by incubation of samples with recombinant PARG (a kind gift from Prof. Dr. Michael O. Hottiger) at a concentration of 1:10,000 (w/w: PARG-to-peptide), at room temperature, overnight and with gentle sample agitation.

### Purification of ADP-ribosylated peptides

After overnight incubation with PARG, samples were cleared from mild precipitation by centrifugation in a swing-out centrifuge at 4°C, for 30 min at 4,250*g*. Next, sepharose beads with GST-tagged Af1521 were added to the samples, in a ratio of 100 μl dry beads per 10 mg sample. Samples were incubated in a head-over-tail mixer, at 4°C for 3 h. Afterwards beads were washed four times in ice-cold IP Buffer, twice in ice-cold PBS, and twice with ice-cold MQ water. Washes were performed in at least 10 bead volumes of washing buffer. After each 2^nd^ washing step, tubes were changed in order to minimize non-specific carryover of contaminants. During the first tube change, after the 2^nd^ wash with IP buffer, all tubes were changed to 1.5 mL Protein LoBind tubes (Eppendorf), and LoBind tubes were used exclusively from this point on throughout the entire purification procedure. Loss of beads was minimized by pelleting beads in a swing-out centrifuge with delayed centrifugation, at 4°C for 3 min at 500*g*. After washing, ADP-ribosylated peptides were eluted off the beads using two bead volumes of ice-cold elution buffer (0.15% TFA). The elution was performed by gentle addition of the elution buffer, gentle mixing of the beads with the buffer every 5 min, and otherwise allowing the beads to stand undisturbed on ice for 20 min. Beads were gently pelleted, and the elutions were transferred to a 0.45 μm column filters (Ultrafree-MC, Millipore), standing on ice. Elution of the beads was repeated once, after which the 2^nd^ elution was pooled with the 1^st^ on top of the 0.45 μm column filter. Co-transfer of beads to the 0.45 um filters was minimized, but not avoided. ADP-ribosylated peptides were then transferred through the filters by centrifugation in a temperature-controlled centrifuge at 4°C, for 1 min at 12,000*g*. After clearance of the samples and removal of any remaining beads, further processing occurred at room temperature. 100 kDa cut-off filters (Vivacon 500, Sartorius) were pre-washed by surface-washing the filters with 300 μL of 50% methanol once, spinning 2× 400 μL of 50% methanol through the filters, surface-washing the filters with 300 μL of 0.15% TFA once, spinning 400 μL of 0.15% TFA through the filters, replacing the collection tubes, and spinning 400 μL of 0.15% TFA through the filters once more, taking care to leave a small amount of liquid on the filters until just prior to usage to prevent their dehydration. Note that this washing procedure of 100 kDa cut-off filters is extremely critical as the filters are preserved in glycerin, of which even trace amounts can obfuscate mass spectrometric analysis and thus ruin the samples. 0.45 μm-cleared ADP-ribosylated peptides were transferred to pre-washed 100 kDa cut-off filters, and centrifuged for 15 min at 8,000*g*, in order to separate the Af1521 macrodomain and other large contaminants from the ADP-ribosylated peptides. Filtered ADP-ribosylated peptides were stored at −80°C until further processing.

### Purification and fractionation of ADP-ribosylated peptides

Initial EThcD energy optimization samples were purified at low pH using StageTips, and all main samples were purified and fractionated at high pH using StageTips. For low-pH (LpH) StageTips, samples were acidified by addition of TFA to 1% (v/v). For high-pH (HpH) StageTips, samples were basified by addition of ammonium hydroxide to a final concentration of 20 mM. All StageTips were prepared essentially as described previously^52^, but were assembled using four layers of C18 disc material (punch-outs from 47mm C18 3M™ extraction discs, Empore). All buffers and samples were passed over StageTips by centrifugation at 1,000*g* until sample loading, and at 1,500*g* during and after sample loading. StageTips were activated using 100 μL methanol, and re-activated using 100 μL of 80% ACN in 0.1% formic acid (for LpH) or using 100 μL of 80% ACN in 50 mM ammonium hydroxide (for HpH). StageTips were equilibrated using 2× 100 μL of 0.1% formic acid (for LpH) or using 2× 100 μL of 20 mM ammonium hydroxide (for HpH), after which samples were loaded. For HpH samples, the flowthroughs were collected at this stage. Subsequently, StageTips were washed with 100 μL of 0.1% formic acid (for LpH) or 100 μL of 20 mM ammonium hydroxide (for HpH). Next, in order to clean the top-end of the StageTips and the top of the C18 plugs, 150 μL of the respective washing buffers were added on top of the C18 plugs, pipetted up and down vigorously while ensuring the buffers reach all the way to the very top of the StageTips, and finally flicked out of the StageTips. This was followed by one final wash with 100 μL of 0.1% formic acid (for LpH) or 100 μL of 20 mM ammonium hydroxide (for HpH). For single-shot LpH samples, elution was performed at this point using 75 μL of 30% ACN in 0.1% formic acid. HpH samples were eluted as fractions, by sequential elution of the StageTips using 75 μL of 2% (F1), 4% (F2), 7% (F3), 10% (F4), 15% (F5), and 25% (F6) of ACN in 20 mM ammonium hydroxide. Flowthroughs from loading of the HpH samples were acidified using 1% TFA (v/v) and subsequently processed at LpH as outlined above, to generate one additional fraction (F0). All samples were vacuum dried to completion in LoBind tubes, using a SpeedVac at 60 °C for 2–3 h. Note that complete drying of the samples is critical for HpH samples, as any residual ammonia will disrupt mass spectrometric analysis of samples. Dried purified ADP-ribosylated peptides were dissolved by addition of 10 μL 0.1% formic acid, allowed to stand without agitation for 5 min, gently tapped to aid final dissolution, briefly centrifuged to move samples to the bottom of the tubes, and finally stored at −20 °C until mass spectrometric measurement.

### Mass spectrometric analysis

All MS samples were measured using a Fusion Lumos Orbitrap mass spectrometer (Thermo). All samples were analyzed on 15-cm long analytical columns, with an internal diameter of 75 μm, and packed in-house using ReproSil-Pur 120 C18-AQ 1.9 μm beads (Dr. Maisch). On-line reversed-phase liquid chromatography to separate peptides was performed using an EASY-nLC 1200 system (Thermo). The analytical column was heated to 40°C using a column oven, and peptides were eluted from the column using a gradient of Buffer A (0.1% formic acid) and Buffer B (80% ACN in 0.1% formic acid). The primary gradient ranged from 3% buffer B to 24% buffer B over 50 minutes, followed by an increase to 40% buffer B over 12 minutes to ensure elution of all peptides, followed by a washing block of 18 minutes. Electrospray ionization (ESI) was achieved using a Nanospray Flex Ion Source (Thermo). Spray voltage was set to 2 kV, capillary temperature to 275°C, and RF level to 30%. Full scans were performed at a resolution of 60,000, with a scan range of 300 to 1,750 m/z, a maximum injection time of 60 ms, and an automatic gain control (AGC) target of 600,000 charges. Precursors were isolated with a width of 1.3 m/z, with an AGC target of 200,000 charges, and precursor fragmentation was accomplished using either electron transfer dissociation (ETD) or electron transfer disassociation with supplemental higher-collisional disassociation (EThcD), using calibrated charge-dependent ETD parameters. For EThcD evaluation, supplemental activation energies of 10, 15, 20, and 25 were used, and a setting of 20 was used for all main experiments. Even though the Orbitrap was exclusively used as the mass detector, both the ion trap and the Orbitrap were calibrated and evaluated in positive and negative mode. For ETD, Reagent Transmission, IC Transmission, and Reagent Ion Source were all calibrated and evaluated on a weekly basis. Calibration of charge-dependent ETD parameters was essentially performed as described previously^53^. Charge-dependent ETD calibration resulted in ETD activation times of 46.42 ms for z=3 precursors, 26.11 ms for z=4 precursors, and 16.71 ms for z=5 precursors. This equates to an ETD Time Constant (τ) of 2.32. Precursors with charge state 3–5 were isolated for MS/MS analysis, and prioritized from charge 3 (highest) to charge 5 (lowest), using the decision tree algorithm. Selected precursors were excluded from repeated sequencing by setting a dynamic exclusion of 48 seconds. MS/MS spectra were measured in the Orbitrap, with a loop count setting of 5, a maximum precursor injection time of 120 ms, and a scan resolution of 60,000.

### Data analysis

Analysis of the mass spectrometry raw data was performed using MaxQuant software (version 1.5.3.30). For both the EThcD energy optimization data search and the main data search, default MaxQuant settings were used, with exceptions outlined below. For generation of the theoretical spectral library, a HUMAN.fasta database was extracted from UniProt on the 10th of July, 2018. N-terminal acetylation, methionine oxidation, cysteine carbamidomethylation, phosphorylation (S, T, and Y), and ADP-ribosylation on all amino acid residues known to potentially be modified (C, D, E, H, K, R, S, T, and Y), were included as variable modifications. For analysis of trypsin samples, up to 6 missed cleavages were allowed. For analysis of Lys-C samples, up to 3 missed cleavages were allowed. A maximum allowance of 4 variable modifications per peptide was used. Second peptide search was enabled (default), and matching between runs was enabled with a match time window of 1 minute and an alignment time window of 20 minutes. Modified peptides were filtered to have an Andromeda score of >40 (default), and a delta score of >20. Data was automatically filtered by posterior error probability to achieve a false discovery rate of <1% (default), at the peptide-spectrum match, the protein assignment, and the site-specific levels.

### Data filtering

Beyond automatic filtering and FDR control as applied by MaxQuant, the data was manually filtered in order to ensure proper identification and localization of ADP-ribose. PSMs modified by more than one ADP-ribose were only rarely observed, and omitted. PSMs corresponding to unique peptides were only used for ADP-ribosylation site assignment if localization probability was >0.90, with localization of >0.75 accepted only for purposes of intensity assignment of further evidences for unique peptides already localized with at least one >0.90 evidence. Because default MaxQuant intensity assignments to modification sites also include non-localized or poorly localized evidences, intensities were manually mapped back to the sites table based on localized PSMs only (>0.90 best-case, >0.75 for further evidences). For the ADP-ribosylation target proteins table, the proteinGroups.txt file generated by MaxQuant was filtered to only contain those proteins with at least one ADP-ribosylation site detected and localized post-filtering as outlined above, with cumulative ADP-ribosylation site intensities based only on localized evidences.

### Comparison to other studies

Data from other studies was retrieved from several publications and online databases. For ADP-ribosylation sites; Larsen et al. (2018), Bonfiglio et al. (2017), Bilan et al. (2017), Leslie Pedrioli et al. (2018). For ADP-ribosylation proteins; Zhang et al. (2013), Jungmichel et al. (2013), Martello et al. (2016), Bilan et al. (2017), Larsen et al. (2018). For total proteome; Bekker-Jensen et al. (2017). For phosphorylation sites; Larsen et al. (2018) and PhosphoSitePlus. For comparison of proteins between studies, all protein identifiers were mapped to the human proteome as downloaded from Uniprot on the 10th of July, 2018. In case data sources did not include Uniprot IDs, ID mapping on Uniprot was used to convert other IDs to Uniprot IDs, and otherwise gene names were used. Non-existent, non-human, or redundant entries, were discarded from the analysis. For comparison of sites between studies, all parent proteins were mapped to Uniprot IDs as described above, and afterwards the reported positions of modification were used to extract 51 amino acid sequence windows (modified residue +/− 25 amino acids), with the sequence windows ultimately used to directly compare identified sites. Sites mapping to non-existent, non-human, or redundant proteins, were discarded. Sites not correctly aligning to the reported amino acid residues were manually corrected in case the modified peptide sequences were available, and otherwise discarded. For generation of the intensity dynamic range plot, all protein intensity values were retrieved from the respective studies, log2 transformed, and the maximum value was subtracted from all other values to have the maxima of all studies align at “0” depth. To minimize interference of data variance at the high end of the intensity scale, standard deviations were calculated over sets of 10 values, and the first value where the SD with the next nine values was <0.2 was considered the maximum intensity value.

### Statistical analysis

Details regarding the statistical analysis can be found in the respective figure legends. Statistical handling of the data was primarily performed using the freely available Perseus software^54^, and includes term enrichment analysis through FDR-controlled Fisher Exact testing. Protein annotations used for term enrichment analysis, including Gene Ontology, UniProt keywords, Pfam, InterPro, and CORUM, were concomitantly downloaded from UniProt with the HUMAN.fasta file used for searching the RAW data. Boxplots were generated using the BoxPlotR web tool^55^. The iceLogo software (version 2.1) was used for sequence motif analysis^56^, and for all sequence comparisons, background sequences were extracted from the same proteins and flanking the same amino acid residue type. Known kinase-regulated sites were extracted from PhosphoSitePlus^41^. The online STRING database (version 10.5) was used for generation of protein interaction networks^57^, and Cytoscape (version 3.7.0) was used for manual annotation and visualization of the STRING networks^58^.

## SUPPLEMENTARY TABLE LEGENDS

**Supplementary Table 1**

A list of all 11,265 unique modified peptides identified with localized ADP-ribosylation sites. Experiment-specific information is included.

**Supplementary Table 2**

A list of all 7,040 localized ADP-ribosylation sites. Fractional modification analysis and experiment-specific information are included.

**Supplementary Table 3**

A list of all 2,149 ADP-ribosylation target proteins. Numbers of ADP-ribosylation sites, site positions, and experiment-specific information are included.

**Supplementary Table 4**

An overview and comparison of 7,681 ADP-ribosylation sites identified across our study and three other ADP-ribosylation proteomics studies.

**Supplementary Table 5**

The complete human proteome, annotated with ADP-ribosylation target proteins identified in this study, as well as in five other ADP-ribosylation proteomics studies. Includes protein copy-numbers as derived from a deep proteome study, and all proteins are annotated with Gene Ontology, Uniprot keywords, Pfam, Interpro, and CORUM terms.

**Supplementary Table 6**

A collection of all statistical information related to term enrichment analyses performed throughout this study.

**Supplementary Table 7**

A list of all 236 unique peptides and 152 sites co-modified by ADP-ribosylation and phosphorylation. Experiment-specific information is included.

